# Variational Autoencoder Modular Bayesian Networks (VAMBN) for Simulation of Heterogeneous Clinical Study Data

**DOI:** 10.1101/760744

**Authors:** Luise Gootjes-Dreesbach, Meemansa Sood, Akrishta Sahay, Martin Hofmann-Apitius, Holger Fröhlich

## Abstract

In the area of Big Data one of the major obstacles for the progress of biomedical research is the existence of data “silos”, because legal and ethical constraints often do not allow for sharing sensitive patient data from clinical studies across institutions. While federated machine learning now allows for building models from scattered data, there is still the need to investigate, mine and understand clinical data that cannot be accessed directly. Simulation of sufficiently realistic virtual patients could be a way to fill this gap.

In this work we propose a new machine learning approach (VAMBN) to learn a generative model of longitudinal clinical study data. VAMBN considers typical key aspects of such data, namely limited sample size coupled with comparable many variables of different numerical scales and statistical properties, and many missing values. We show that with VAMBN we can simulate virtual patients in a sufficiently realistic manner while making theoretical guarantees on data privacy. In addition, VAMBN allows for simulating counterfactual scenarios. Hence, VAMBN could facilitate data sharing as well as design of clinical trials.

## Introduction

Clinical studies are important to increasingly base medical decisions on statistical evidence rather than on personal experience. Within a given area of disease there can exist many studies, and each of these studies has unavoidably certain biases due to inclusion / exclusion criteria or over-representation of specific geographic regions and ethnicities. Moreover, usually neither the same clinical outcome measures nor the same molecular data are systematically collected in different studies of the same disease. Accordingly, compilation of a comprehensive view of a specific disease requires to analyze and compare multiple studies. However, legal and ethical constraints typically do not allow for sharing sensitive patient data beyond summary statistics outside the organization that is the owner, and even within one and the same organization the same reasons sometimes prevent data sharing. In consequence there exist data “silos”. This is increasingly becoming an issue as medicine as a whole is becoming more and more driven by the availability of Big Data and their analysis, including the increasing use of Artificial Intelligence (AI) and in particular machine learning methods in precision medicine (Fröhlich et al., 2018). While recent developments of federated machine learning techniques are certainly a major step forward (Ghosh et al., 2019; McMahan et al., 2016), their implementation is technically challenging and does not permit researchers to unbiasedly investigate, mine and understand clinical study data located within different organizations.

Sufficiently realistic simulations of virtual patient cohorts could not not only be a mechanism to break data “silos”, but also to allow researchers to conduct counterfactual experiments with patients, e.g. in the context of intensive care units (Chase et al., 2018; Knab et al., 2016) or for better design of clinical trials (Galbusera et al., 2018; Lim et al., 2017). Regarding the latter we should mention that most existing work on virtual trial simulation focuses on modeling of mechanistically well understood pharmacokinetic and pharmacodynamic processes (Holford et al., 2010; Pappalardo et al., 2018). In contrast, our focus is here on data driven, model based simulations of virtual patients across biological scales and modalities (e.g. clinical, imaging) where no or little mechanistic understanding is available and required.

We suggest a generative modeling framework for simulation of virtual patients, which is specifically designed to address the following key features of clinical study data:

- limited sample size in the order of a few hundred patients
- highly heterogeneous data with many variables of different distributions and numerical scales
- longitudinal data with many missing values

Our novel proposed method (Variational Autoencoder Modular Bayesian Networks - VAMBN) is a combination of a Bayesian Network (Heckerman, 1997) with modular architecture and Variational Autoencoders (Kingma and Welling, 2013) encoding defined groups of features in the data. Due to its specific design VAMBN does not only allow for generating virtual patients under certain theoretical guarantees for data privacy (Dwork et al., 2006a), but also for simulating counterfactual interventions within them, e.g. a shift by age. Moreover, we demonstrate that one can “learn” the conditional distribution of a feature in one study to counterfactually add it to another one.

We evaluate our VAMBN on the basis of two Parkinson’s Disease (PD) studies, where we show that marginal distributions, correlation structure as well as expected effects (treatment effect on motor symptoms and difference of clinical outcome measures to healthy controls, respectively) are largely preserved in simulated patients. Moreover, we demonstrate that counterfactual simulation results match general expectations. Finally, we show that VAMBN models capture expected causal relationships in the data.

## Methods

### Motivation and Conceptual Idea Behind VAMBN

Our proposed approach rests on the idea of learning a generative model of longitudinal clinical study data. It combines two classes of generative modeling techniques: Bayesian Networks (BNs) (Heckerman, 1997) and Variational Autoencoders (VAEs) (Kingma and Welling, 2013). Bayesian Networks (BNs) are probabilistic graphical models, which represent a joint statistical distribution of several random variables by factorizing it according to a given directed acyclic graph into local conditional statistical distributions. Attractive properties of BNs are

- efficient encoding of multivariate distributions
- interpretability, because the graph structure can be used to represent causal relationships
- a theoretical framework to simulate interventions via the “do” calculus (Pearl, 2000)

Unfortunately, under general conditions inference within a BN and learning of the graph structure from data are both NP-hard computational problems (Koller and Friedman, 2009). Computationally efficient parameter and structure learning can only be achieved, if all random variables follow multinomial or Gaussian distributions. However, this scenario is in reality too restrictive for many applications, including clinical study data, where many variables do not follow any known parametric distribution. In addition, the NP hardness of BN structure learning raises severe concerns, because clinical study data has often dozens of variables (measured over time). But, the number of patients is typically only in the order of a few hundred. Hence, the chance to identify the correct graph from these limited data is questionable.

VAEs are a neural network based approach that maps input data to a low dimensional latent distribution (typically a Gaussian) through several sequential encoding steps. VAEs are typically trained via stochastic gradient descent to optimize an evidence / variational lower bound (ELBO) on the log-likelihood of the data. VAEs have recently been extended to deal with heterogeneous multi-modal and missing data (Nazabal et al., 2018), which is the common situation in clinical studies. VAEs are generative, because drawings from the latent distribution can be decoded again. A limitation of VAEs is that in a situation with comparably small data a dense VAE model with several hidden layers could easily overfit. Moreover, interpretation of the neural network models is far more challenging than for BNs.

Our suggested approach aims for combining the advantages of BNs and VAEs while mitigating their limitations (Figure 1): Following the idea of Module Networks (Segal et al., 2003, 2004) we first define modules of variables that group together according the design of the study. For example, demographic features, clinical assessment scores, medical history, treatment might each form such a module. Our aim is then to learn a BN between these modules. In contrast to Segal et al. we do not use regression trees to represent conditional joint distributions of variables within each module, but VAEs, because they are generative. Each VAE is thus only trained on a small subset of variables, hence significantly reducing the number of network weights compared to a full VAE model for the entire dataset and allowing for applying the well established “do” calculus for simulating interventions. We call our approach Variational Autoencoder Modular Bayesian Network (VAMBN). Due to its generative nature VAMBN allows for simulating virtual subjects by first drawing a sample from the BN and second by decoding it through the VAE representing the corresponding module.

We validate virtual patient cohorts by comparing against original patients

- marginal distributions of individual variables
- correlation structures
- expected differences between patient subgroups, e.g. treated vs placebo patients

In the following we explain the individual steps of our method in more detail and we discuss, how data privacy can be theoretically guaranteed.

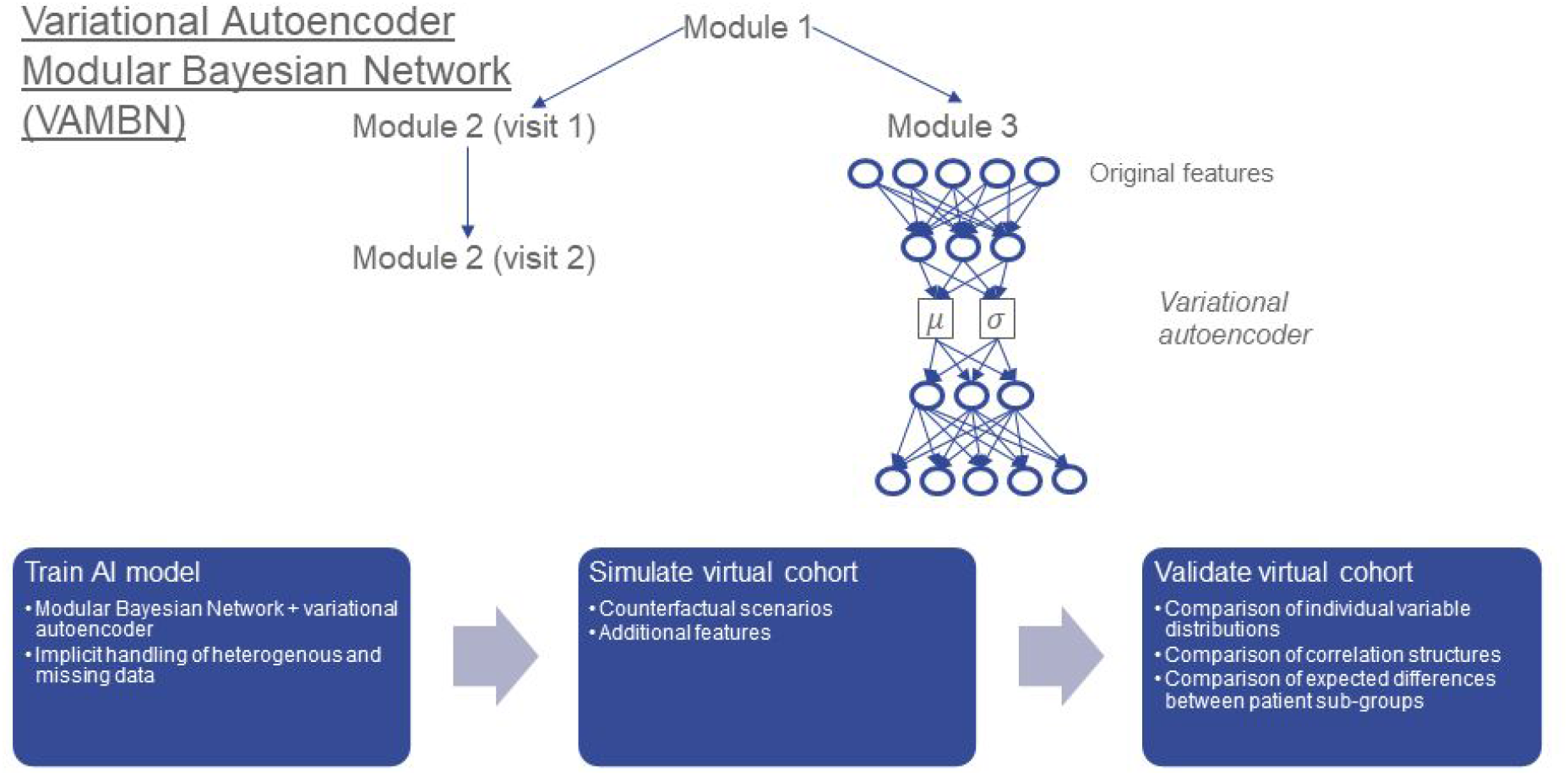

### Modular Bayesian Networks (MBNs)

The starting point of our proposed approach is a Modular Bayesian Network (MBN) describing in a longitudinal manner statistical dependencies of VAE encoded modules (i.e. sets of variables in the original data): Let *G* = (*V, E*) be a directed acyclic graph (DAG) and *X* = (*X*_*υ*_)_*υ*∈*V*_ a set of random variables indexed by nodes in *V. X* is called a Bayesian Network (BN) with respect to *G*, if the joint distribution *p*(*X*_1_, *X*_2_, … *X*_*n*_) factorizes according to:

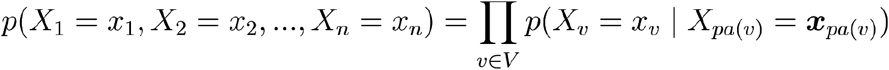

where *pa*(*υ*) denotes the parents of node *υ* and ***x***_*pa*(*υ*)_ their joint configuration (Koller and Friedman, 2009). For a given node *υ* we summarize the set of associated conditional probabilities into a parameter vector *θ*_*υ*_, and these parameter vectors are assumed to be statistically independent for different nodes *υ*, *υ*′.

In our situation there exists a subset 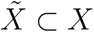 that is time dependent, i.e. 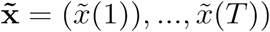 with *T* being the number of visits. Dynamic Bayesian Networks (Ghahramani, 1998) usually deal with this situation by implicitly unfolding the BN structure over time, i.e. introducing for each visit *t* a separate copy 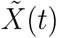 of 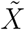 while requiring that edges always point from time slice *t* to time slice *t* + 1 (corresponding to a first order Markov process). This implicit unfolding assumes a stationary Markov process, i.e. parameters *θ* do not change with time. In our setting this assumption is most likely wrong, because patients change in their disease outcome during the course of a study, i.e. 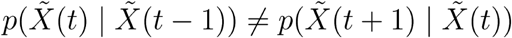. Hence, we here use an unfolding strategy, in which we explicitly use different copies 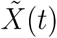 for each time point. In addition, unfolding of the BN structure saves us from modeling the dynamical behavior of the data within the VAE framework (e.g. via LSTM units - (Hochreiter and Schmidhuber, 1997)), which would require far more parameters.

In our case nodes (i.e. random variable) either follow a Gaussian distribution (we explain the reasons later), or they could be of categorical nature, i.e. follow a multinomial distribution and not be autoencoded. A restriction we impose at this point is that a discrete node cannot be the child of a Gaussian one. Under this assumption the conditional log-likelihood of the training data et al., 2018): *D* = {*x*_*vi*_ | *i* = 1, …, *N*, *v* ∈ *V*} given *G* can be calculated analytically (Andrews et al., 2018):

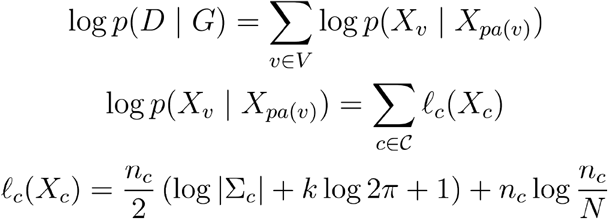

where *𝓒* is the set of possible partitionings of Gaussian variable *X*_*υ*_ according to the configuration of its discrete parents, and *n*_*c*_ is the number of patients in partition *c*. *X*_*c*_ is the associated design matrix, and *k* the number of columns of that matrix. In a similar way the local log-likelihood *ℓ*_*c*_(*X*_*c*_) for a discrete node *X*_*υ*_ with only discrete parents can be computed. By considering in addition the number of parameters of the MBN we can use the Bayesian Information Criterion (BIC) to score *G* with respect to data *D*. In practice we make use of the corresponding implementation in R-package bnlearn (Scutari, 2010).

### Modeling Missing Data in MBNs

One of the key challenges with longitudinal patient data is missing values, which can result due to different reasons: a) patients drop out of a study, e.g. due to worsening of symptoms; b) a certain diagnostic test is not taken at a particular visit (e.g. due to lack of patient agreement), potentially resulting in missing information for entire variable groups; c) unclear further reasons, e.g. time constraints, data quality issues, etc. From a statistical point of view these reasons manifest into different mechanisms of missing data (Kang, 2013; Rubin, 1976):

- missing completely at random (MCAR): The probability of missing information is not related to either the specific value which is supposed to be obtained or other observed data. Hence, entire patient records could be skipped without introducing any bias. However, this type of missing data mechanism is probably rare in clinical studies.
- missing at random (MAR): The probability of missing information depends on other observed data, but is not related to the specific missing value which is expected to be obtained. An example would be patient drop out due to worsening of certain symptoms, which are at the same time recorded during the study.
- missing not at random (MNAR): any reason for missing data, which is neither MCAR or MAR. MNAR is problematic, because the only way to obtain unbiased estimates is to model missing data.

Missing values in clinical study data are most likely a combination of MAR and MNAR mechanisms. In general, multiple imputation methods have been proposed to deal with missing data in longitudinal patient data (Kang, 2013). Specifically for MNAR it has been suggested to explicitly encode systematic missingness of variables or variable groups via dedicated indicator variables (Mustillo and Kwon, 2015). The missing value itself can technically then be filled in by any arbitrary value, e.g. zero.

In our MBN framework auxiliary variables are fixed parents of all nodes, which contain missing values in a non-random way. There also exist higher level missing data nodes that show whether a participant does not have any data for the entire visit. If the auxiliary variable of a node representing an autoencoded variable group is identical to the missing visit node, the auxiliary variable itself is removed from the network and the node is directly connected to the missing visit node instead. These higher level nodes account for the high correlation between the different auxiliary nodes at a visit. Note that to facilitate modelling in the MBN, auxiliary and missing visit nodes were only introduced for nodes and visits with more than 5 missing data points in total.

### MBN Structure and Parameter Learning

#### Structure Learning

Most edges in the MBN structure are not known and hence need to be deduced from data. Unfortunately, MBN structure learning is an NP hard problem, because the number of possible DAGs grows super-exponentially with the number of nodes (Chickering et al., 2004). Hence, the search space of possible network structures should a priori be restricted as much as possible. We follow two essential strategies for this purpose:

1. We group variables in the raw data into autoencoded modules, as explained above.
2. We impose causal constraints on possible edges between modules.

More specifically, we imposed the following type of constraints:

- Modules of demographic and other clinical baseline features (e.g. age, gender, ethnicity) can only have outgoing edges..
- Modules representing medical history can only depend on the modules mentioned in 1. and biomarkers.
- Modules of imaging features can be related to each other, but they don’t influence other modules.
- Modules of clinical outcome measures (e.g. UPDRS) can influence imaging and they can be mutually correlated with assessment of non-motor symptoms.
- Biomarker modules can influence all modules, except for modules of clinical baseline features.
- Longitudinal measures must follow the right temporal order, i.e. there are no edges pointing backwards in time.
- Empirically proven edges (e.g. the treatment effect on the first maintenance visit in SP513 data) must be reflected in the network structure.
- Auxiliary and missing visit nodes were connected to their respective counterparts at the next time point, accounting for a correlation between these measures over time, e.g. through study dropout.

Accordingly, we blacklisted possible edges that could violate any of these constraints. Structure learning was then conducted via tabu search (Hong et al., 2016), which is essentially a modified hill climber that is designed to better escape local optima. This choice was made, because score based search algorithms have empirically found to show a more robust behavior in terms of network reconstruction accuracy than constraint based methods for mixed discrete / continuous data, specifically for smaller sample sizes (Raghu et al., 2018). In addition, it should be noted that due to the typical small number variables in the MBN runtime was not a major concern here.

#### Parameter Learning

Given a graph structure *G* of a MBN parameters (i.e. conditional probability tables and conditional densities) can be estimated via maximum likelihood. Note that estimation of the conditional Gaussian density for a node *V* amounts to fitting a linear regression function with parents of *V* being predictor variables. Conditional probability tables, on the other hand, can be estimated by counting relative frequencies of *V* taking on a particular value *υ*.

### Variational Autoencoders

VAEs were introduced by (Kingma and Welling, 2013) and can be interpreted as a special type of Bayesian Network, which has the form *Z* → *X*, where *Z* is a latent, usually multivariate standard Gaussian and *X* a multivariate random variable describing the input data. Moreover, for any sample (*x*, *z*) we have *p*(*x* | *z*) = *N* (μ(*z*), σ(*z*)). One of the key ideas behind VAEs is to variationally approximate

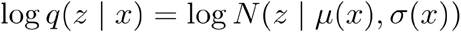

That means μ(*x*) and σ(*x*) are the mean and standard deviation of the approximate posterior and are outputs of a multilayer perceptron neural network that is trained to minimize for each data point *x* the ELBO criterion

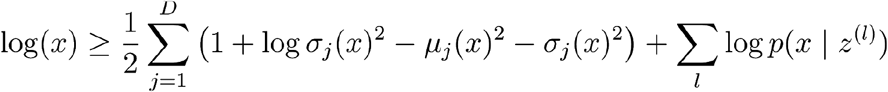

where *z* = *μ*(*x*) + *σ*(*x*) ⊙ *ϵ*^(*l*)^ with *ϵ*^(*l*)^ ~ *N*(0, *I*). Here ⊙ denotes an element-wise multiplication.

### Variational Autoencoders for Heterogeneous and Incomplete Data (HI-VAE)

VAEs were originally developed for homogenous data without missing values. But, clinical data within one and the same module (e.g. demographics) could contain continuous as well as discrete features of various distributions and numerical ranges, i.e. the data is highly heterogeneous. Moreover, there could be missing values. Recently, (Nazabal et al., 2018) extended VAEs to address this situation. Their HI-VAE approach starts from a factorization of the VAE decoder according to

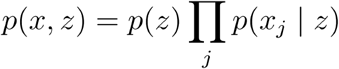

where *x* ∈ ℝ^*D*^ denotes a D dimensional data vector and *z* ∈ ℝ^*K*^ its *K* dimensional latent representation. Furthermore, *x*_*j*_ indicates the *j*-th feature in *x*. In the factorization it is further possible to separate observed (*𝓞*) from missing features (*𝓜*):

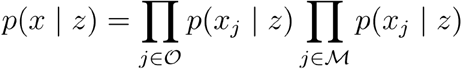

A similar separation is possible in the decoder step. Accordingly, VAE network weights can be optimized by solely considering observed data (input drop-out model). Note that the input drop-out model is essentially identical to the approach we described earlier for MBNs.

To account for heterogeneous data types Nazabal et al. suggest to set

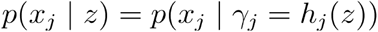

where *h*_*j*_(·) is a function learned by the neural network, and *γ*_*j*_ accordingly models data modality specific parameters (e.g. for real-valued data 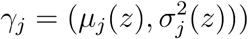. Moreover, the authors use batch normalization to account for differences in numerical ranges between different data modalities. Finally, Nazabal et al. do not use a single Gaussian distribution as a prior for *z*, but a mixture of Gaussians, i.e.:

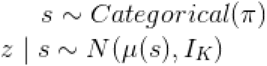

where *s* is *K* dimensional. We refer to (Nazabal et al., 2018) for more details about their VAE extension. Importantly, categorical variables *s* are added to the MBN graph *G* as parents of variables encoding modules. In practice we kept *K* at 1 for all modules, resulting in a single normal distribution for *z*, with the exception of the demographic data in both studies and the neurological examination in SP513 data. For these modules *K* was set to 2, as training suggested that a single normal distribution was not a good fit for the data.

### VAMBN: Bringing MBNs and HI-VAEs Together

Let *Y*_*υ*_ denote the set of observed variables associated to module *Y*_*υ*_ denote the set of observed variables associated to module *X*_*υ*_, *υ* ∈ *V*. Note that variables *X*_*υ*_ are low dimensional embeddings / encodings of the *Y*_*υ*_. The total likelihood *p*(*X, Y* | *G*, Θ) given graph *G* and model parameters Θ can be written as:

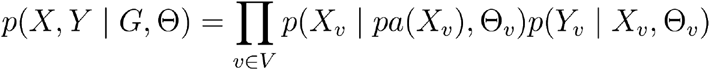

where *p*(*Y*_*υ*_ | *X*_*υ*_, Θ_*υ*_) is the generative model of the data represented by HI-VAE (it is the decoder distribution). Moreover, *pa*(*X*_*υ*_) denotes all module nodes plus (in our case 1 dimensional) categorical *s* variables, see last Section. Hence, *p*(*X*_*υ*_ | *pa*(*X*_*υ*_), Θ_υ_) is a normal distribution with mean

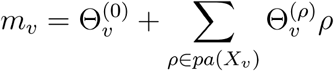

(i.e. modeled via a linear regression with intercept 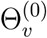 and slope coefficients 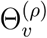), and residual variance *ν*_*υ*_ = *Var*(*X*_*υ*_ − *m*_*υ*_).

Our aim is to find parameters Θ maximizing log *p*(*X, Y* | *G*, Θ). Using the factorization of this quantity and the typical assumption of node-wise statistical independence of parameters (Koller and Friedman, 2009) we can optimize the total log-likelihood by the following two steps:

1. For all 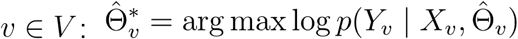. This is achieved via training a HI-VAE model for each module, i.e. optimizing associated network weights 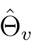.
2. For all 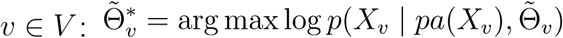. This is achieved by learning the MBN structure *G* and associated parameters 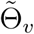 based on HI-VAE encoded modules.

Overall, the training of the proposed VAMBN approach thus consists of the following steps:

1. Definition of modules of variable.
2. Training of HI-VAEs for each module. In practice the training procedure included a hyper-parameter optimization over

a. learning rate ∈ {0.01, 0.001}
b. mini-batch size ∈ {16, 32} Each candidate parameter set was evaluated via a 3-fold cross-validation using the reconstruction loss as objective function.
3. Definition of constraints for possible edges in the MBN.
4. Structure and parameter learning of the MBN using encoded values for each module: Note that by construction of our model each variable *X*_*υ*_ follows a mixture of Gaussian distributions. Let *s* ~ *Categorical*(*π*) indicate the mixture component. Hence, *X*_*v*_ | *s* is Gaussian. Introducing *s* into the MBN thus yields a network with only Gaussian and discrete nodes, and parameter and structure learning can accordingly performed computationally efficiently, as explained before.

We also considered to use *N*(*m*_*υ*_, *ν*_*υ*_) as a prior for *X*_*υ*_ instead of the original Gaussian mixture prior for training of HI-VAE models in a second iteration of the entire VAMBN training procedure. In reality we could not observe a significant increase of the total model likelihood *p*(*X, Y* | *G*, Θ) due to this computationally more costly procedure, see section A of the Supplementary Materials. Reported results hence only refer to the original VAMBN approach without any further continued training using a modified prior.

### Simulating Virtual Patients and Counterfactual Scenarios

The trained VAMBN model can be used to create a virtual patient cohort. Virtual patients are simulated as follows:

1. Draw samples from the MBN. This can be achieved by following the topological order of nodes in the DAG. That means we first sample from the conditional distribution of parent nodes, before we do the same for their children while conditioning on the already drawn values each of the parents.
2. Decode MBN samples through HI-VAE. Note that a sample drawn from the MBN represents a vector of latent codes. Decoding maps these codes back into the original input space.

To perform a simulation of a counterfactual situation we rely on the ideal intervention scheme established by Pearl (Pearl, 2000). That means rather than sampling from a joint distribution *p*(*X*_1_, *X*_2_, …, *X*_*n*_) we draw from *p*(*X*_1_, *X*_2_, …, *X*_*p*−1_, *X*_*p*+1_, …, *X*_*n*_ | *do*(*X*_*p*_ = *x*)) where *X*_*p*_ denotes a variable in the MBN, in which we simulate the counterfactual scenario *X*_*p*_ = *x*. Practically this can be achieved by deleting all incoming edges to *X*_*p*_ in the MBN structure, setting *X*_*p*_ = *x* and then drawing from the modified MBN. Subsequently, the variables can be decoded through the HI-VAE, as described before.

### Using VAMBN for Counterfactually Adding Features to a Dataset

A special case of the counterfactual simulation described in the last Section is the addition of features to a dataset, which have not been observed within a particular study A, but within another study B: Let *Y* be a (module of) variables in study B not observed in A. We assume the existence of MBNs 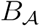 and 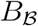 for both datasets. Moreover, we suppose 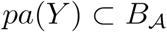, i.e. parents of *Y* are also in A. Hence, we can draw from the interventional distribution

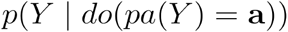

where a denotes a configuration of parent nodes of *Y* observed in dataset A. Therefore, we can counterfactually add for any patient in dataset A possible values for *Y* by considering his/her observed features that may impact *Y*.

### Differential Privacy Respecting Model Training

One of our motivations for developing VAMBN was to enable a mechanism for sharing data across organizations that addresses data privacy concerns. Practically this could be achieved by sharing either simulated datasets or ready trained VAMBN models. However, specifically in the latter case there is the concern that by systematically feeding inputs and observing corresponding model outputs it might be possible to re-identify patients that were used to train VAMBN models. This is particularly true for HI-VAEs, which encode groups of raw features.

Differential privacy is a concept developed in cryptography that poses guarantees on the probability to compromise a person’s privacy by a release of aggregate statistics from a dataset (Dwork et al., 2006b): Let *𝒜* be a randomized algorithm and 0 < *ϵ*, 0 < *δ* < 1. According to (Dwork et al., 2006a) 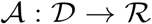 is said to respect (*ϵ*, *δ*) differential privacy, if for any two datasets *D*_1_, *D*_2_ ∈ *𝒟* that differ only in one single patient and for any output of the randomized algorithm 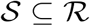 we have

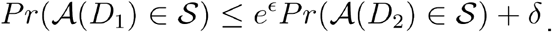

(Abadi et al., 2016) showed that it is possible to directly incorporate (*ϵ*, *δ*) differential privacy guarantees into the training of a neural network by clipping the norm of the gradient and adding a defined amount of noise to it.

It is straightforward to incorporate this approach into the training of each of the VAE models within VAMBN. Hence we are able to provide guarantees on (*ϵ*, *δ*) differential privacy for the entire VAMBN model, because (*ϵ*, *δ*) differential privacy is composable. That means the property for system of several components is fulfilled, if all of its components fulfill (*ϵ*, *δ*) differential privacy (Dwork et al., 2006a).

## Data

### SP513

SP513 was a randomized, double blinded and placebo controlled study to compare two PD drugs within an early disease population (Giladi et al., 2007). We here examine 557 patients of the final analysis set, which had received treatment. Out of these patients 117 received placebo, 227 ropinirole and 213 another dopamine agonist. Both drugs were first up-titrated within a 13 weeks time period and then followed up for 24 weeks. We model the screening and baseline visits as well as three visits in the maintenance phase. Clinical variables captured during the trial comprised baseline demographics, disease duration, UPDRS scores, Epworth Sleepiness Scale (ESS), Hoehn & Yahr stage and standard blood biomarkers for safety assessment (e.g. hemoglobin, creatinine, etc.).

### PPMI

The Parkinson’s Progression Markers Initiative (PPMI) (www.ppmi-info.org/data) consists of multiple cohorts from a network of clinical sites with the aim to identify and verify progression markers in PD. It is a multi-modal, longitudinal observation study with data collected using standardized protocols (Parkinson Progression Marker Initiative, 2011). PPMI comprises of eight cohorts with different clinical and genetic characteristics. Here we used data of 362 de novo PD patients and 198 healthy controls. All PD patients were initially untreated and diagnosed with the disease for two years or less. They showed signs of resting tremor, bradykinesia, and rigidity. We used 266 clinical variables measured at 11 visits during 96 months comprising demographics, patient PD history, DaTSCAN imaging, non-motor symptoms, CSF (cerebrospinal fluid) biomarkers (A- β, α -synuclein, dopamine, phospho-tau, total tau) and UPDRS scores.

## Results

### VAMBN Reflects Expected Causal Relationships in Data

As outlined in the Methods part of this paper our proposed VAMBN approach results into a Modular Bayesian Network that describes conditional statistical dependencies between groups of variables that are encoded via HI-VAEs. An obvious initial question is whether learned dependencies between modules reflect expected causal relationships and, if yes, how statistically stable these can be detected. To address this point we performed a non-parametric bootstrap of the MBN structure learning (Davison and Hinkley, 1997). That means that for each study, we resampled the existing *N* patients 1000 times with replacement. For each bootstrap dataset we ran a complete MBN structure learning, and we counted the fraction of times that each edge was included in the model. We overlayed this bootstrapped network with the MBN learned from the complete data to get an overall impression of the learned VAMBN model as well as the stability of inferred conditional statistical dependencies.

Figures 2a), b) highlight that in both SP513 as well as PPMI inferred edges agree well with expected causal dependencies: For example, in SP513 (Figure 2a) UPDRS scores of subsequent visits are connected with each other, and impact sleepiness (ESS). ESS itself is dependent on medical history. UPDRS scores are during the titration phase influenced by Hoehn & Yahr stages and the illness severity score defined in SP513. Safety biomarkers depend on gender, but otherwise have no impact.

**Figure 2.**
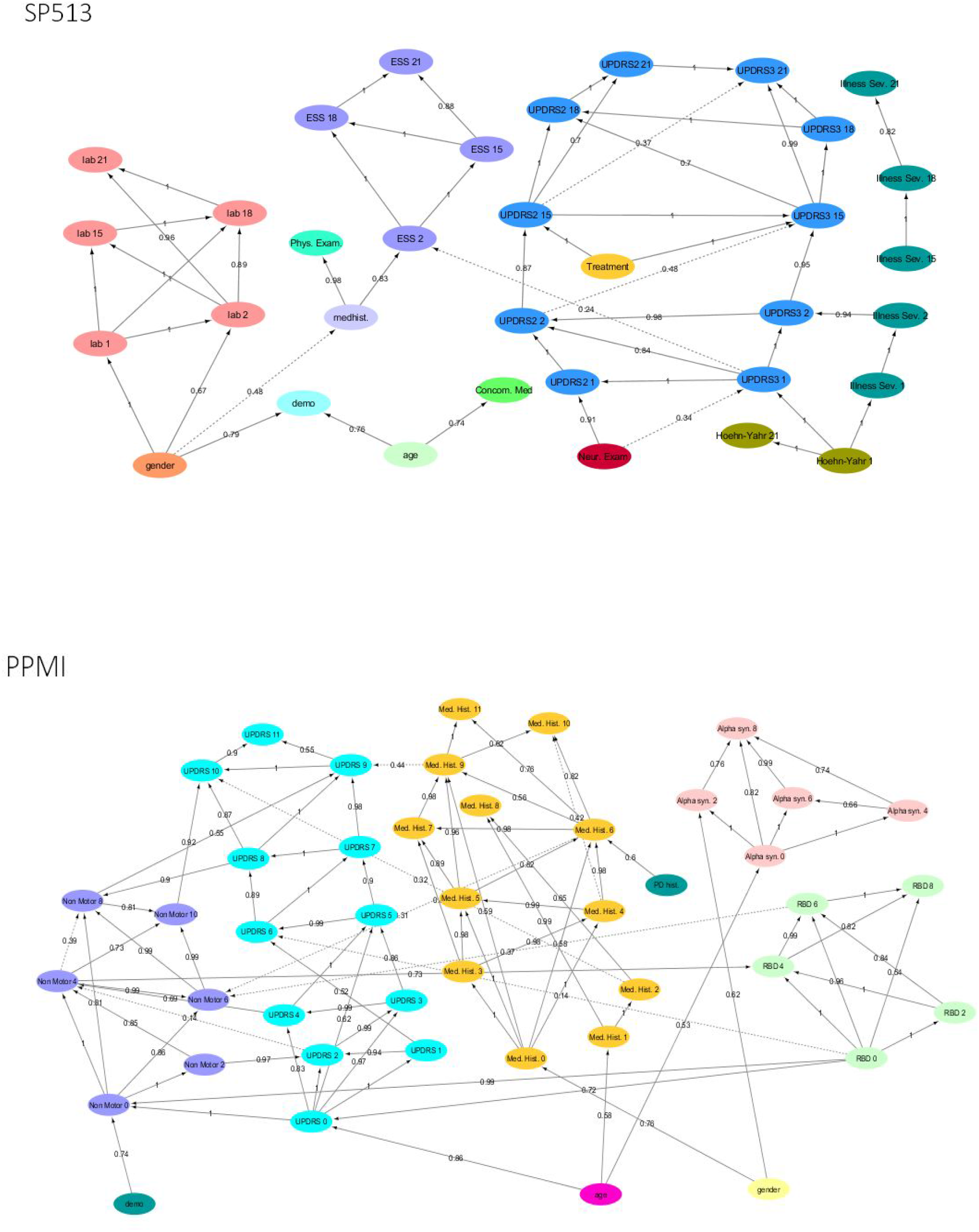
Final MBNs learned by VAMBN based on SP513 and PPMI data. The edges are labelled with the bootstrap frequencies of each connection. For readability, auxiliary variables and missing visit nodes were removed for the visualisation. Figures are also available as Cytoscape files in the Supplements for better convenience.

In PPMI (Figure 2b) the RBD sleepiness score and non-motor symptoms mutually influence each other, and the same holds true for UPDRS. UPDRS is dependent on age, medical history and α -synuclein levels in CSF.

Altogether these examples underline that VAMBN models permit a certain level of interpretation.

### Simulated Patients are Realistic

Simulated patient trajectories generated by VAMBN are only useful, if they are sufficiently similar to real ones. On the other hand we clearly do not want VAMBN to simply re-generate the data it was trained on (which would trivially maximize similarity to real patients). It is therefore not straightforward to come up with a criterion or interpretable index to measure the quality of a virtual patient simulation.

From our point of view simulated patients should mainly fulfill three criteria:

a. Summary statistics (e.g. mean, variance, median, lower quartile, upper quartile) over individual variables should look similar to real ones.
b. Correlations between variables in simulated patients should be close to the ones observed in real ones.
c. Treatment effects or other expected outcomes should be similar in simulations, also in terms of effect size.

To assess VAMBN with respect to these criteria we simulated the same number of virtual patients as real ones in each of the two PD studies. Figure 3 demonstrates that marginal distributions for individual variables were in general sufficiently similar (but not identical) to the empirical distributions of real data in both PD studies. For additional plots see section B of the Supplementary Materials. In addition, the empirical distributions of Pearson correlations in simulated and real data were close to each other (Figure 4). Interestingly, in both cases (marginal distributions and correlations) largest differences were observed between HI-VAE decoded features of real patients and original features of the same patients. Hence, the majority of the “simulation error” can be attributed to an imperfect fit of HI-VAE models.

**Figure 3.**
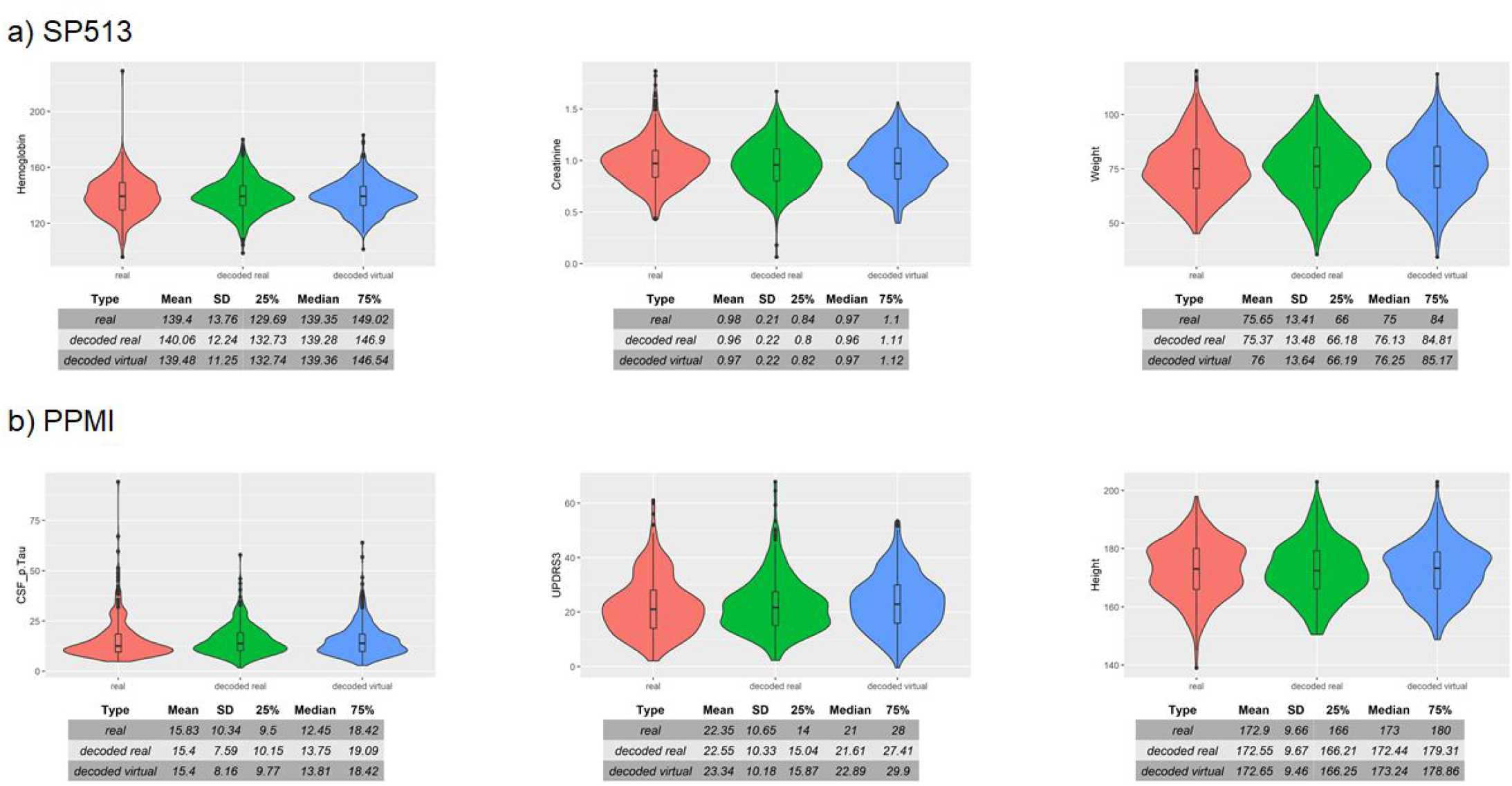
Examples of real and simulated / virtual patients for SP513 (a) and PPMI (b) datasets. The Figure compares the marginal distributions of selected variables for real patients (red), virtual patients (blue) and real patients decoded via the HI-VAE model (green). Tables show summary statistics of the distributions. Plots and tables of further variables can be found in section B of the Supplementary Materials.

**Figure 4.**
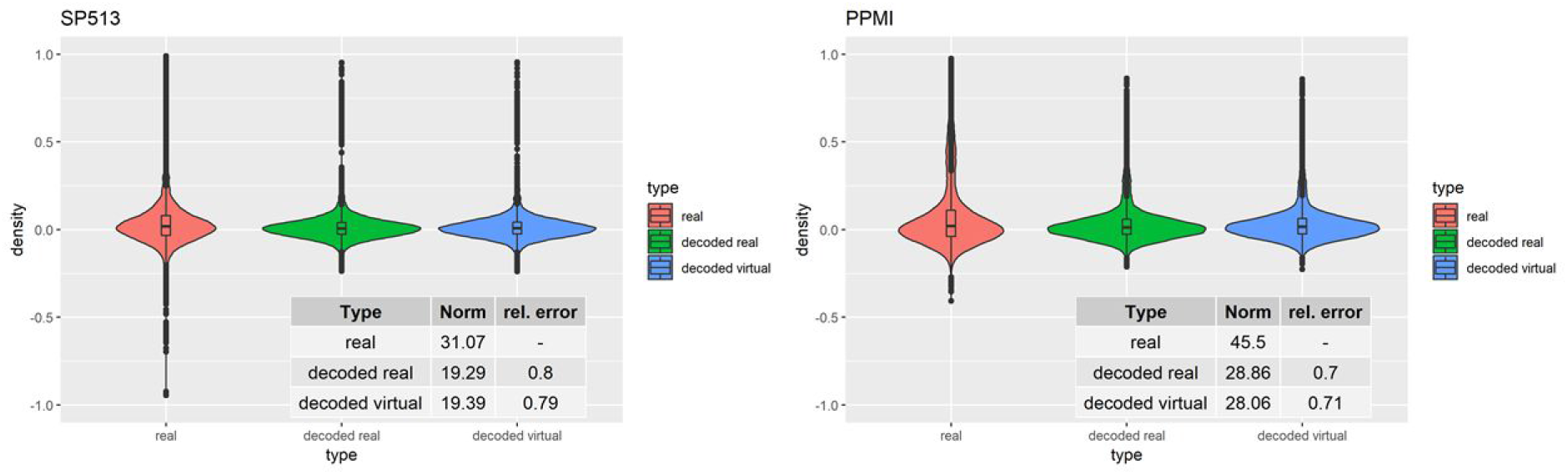
Distribution of Pearson correlation coefficients between variables in real patients (red), virtual patients (blue) and decoded real patients (green). Tables show the Frobenius norm of the correlation matrices as well as the relative error, which consists of the norm of the matrix that is the difference between the decoded real or virtual correlation matrix divided by the norm of the original correlation matrix.

UPDRS3 scores in simulated PD patients showed similar differences to healthy controls than real PD patients im PPMI (Figure 5 right). Moreover, the ropinirole treatment effect in simulated and real SP513 patients demonstrated a comparable effect size and p-value (Figure 5 left).

**Figure 5.**
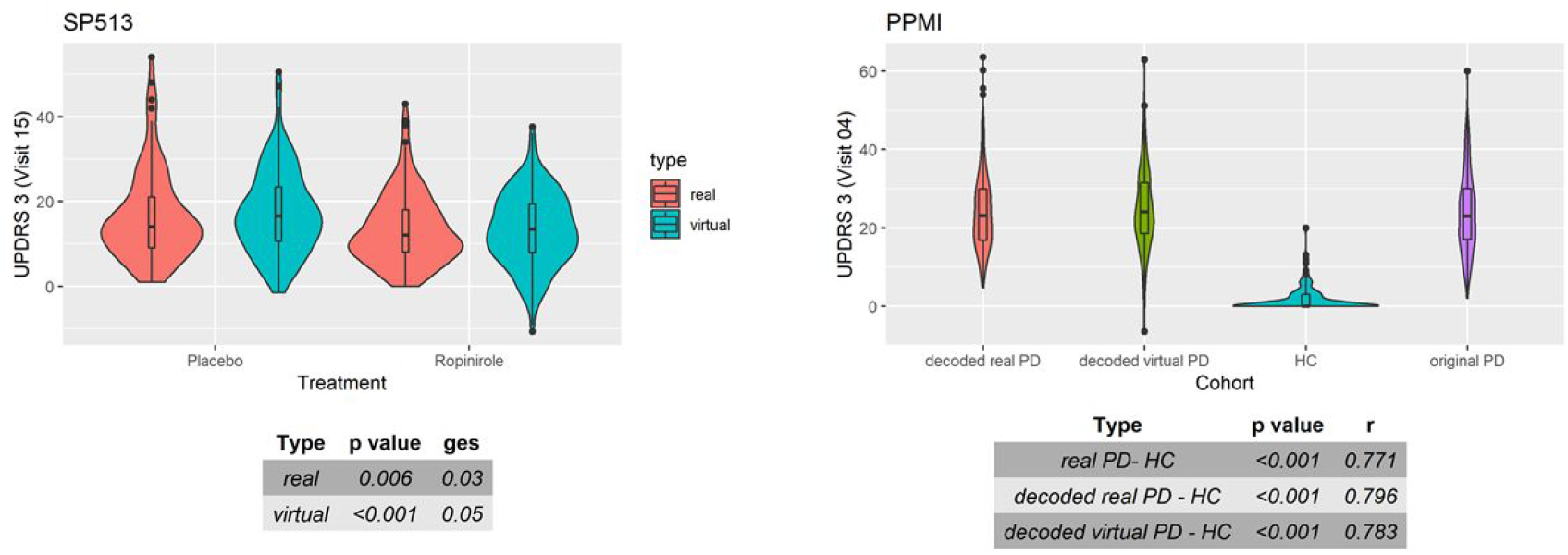
Distribution of UPDRS3 scores in SP513 (left) and PPMI (right). Left: The plot depicts in red UPDRS3 scores of real SP513 patients under placebo and ropinirole at visit 15 (i.e. during treatment phase), respectively. In blue the distribution of the UPDRS3 score in the same number of virtual patients is shown. Effect sizes and corresponding p-values obtained from two one-way ANOVAs comparing placebo and drug treatment in the real and virtual patients are shown in the tables at the bottom. Similar plots at other visits can be found in section C of the Supplementary Materials. Right: Distribution of original (purple) and decoded (red) UPDRS3 scores of real PPMI de novo PD patients at visit 4 in comparison to PPMI healthy controls (blue). UPDRS3 scores of virtual PD patients are shown in yellow. The table at the bottom shows differences in UPDRS3 scores between original PD, decoded real PD and virtual PD patients compared to PPMI healthy controls, showing p-value and effect size from three Mann-Whitney U tests. Similar plots at other visits can be found in section C of the Supplementary Materials.

Altogether we thus concluded that VAMBN allows for a sufficiently realistic simulation of virtual subjects with respect to our three defined criteria. At the same time we could confirm that indeed none of the simulated patients was a simple re-generation of one of the patients in the training data.

### Generalizability of VAMBN Models

A relevant question is, how generalizable VAMBN models are, i.e. whether they are purely overfitted or whether they can sufficiently describe data in an independent test set. To address this point we randomly split data in SP513 and PPMI into 80% training and 20% test. VAMBN models were only fitted to the training set. We then recorded the log-likelihood of patients in the training set and test set, indicating a sufficiently good agreement (Figure 6). We thus concluded that VAMBN models are generally not overfitted. That means the previously reported agreement of virtual and real patients cannot just be the result of overfitting the data with an overly complex model.

**Figure 6.**
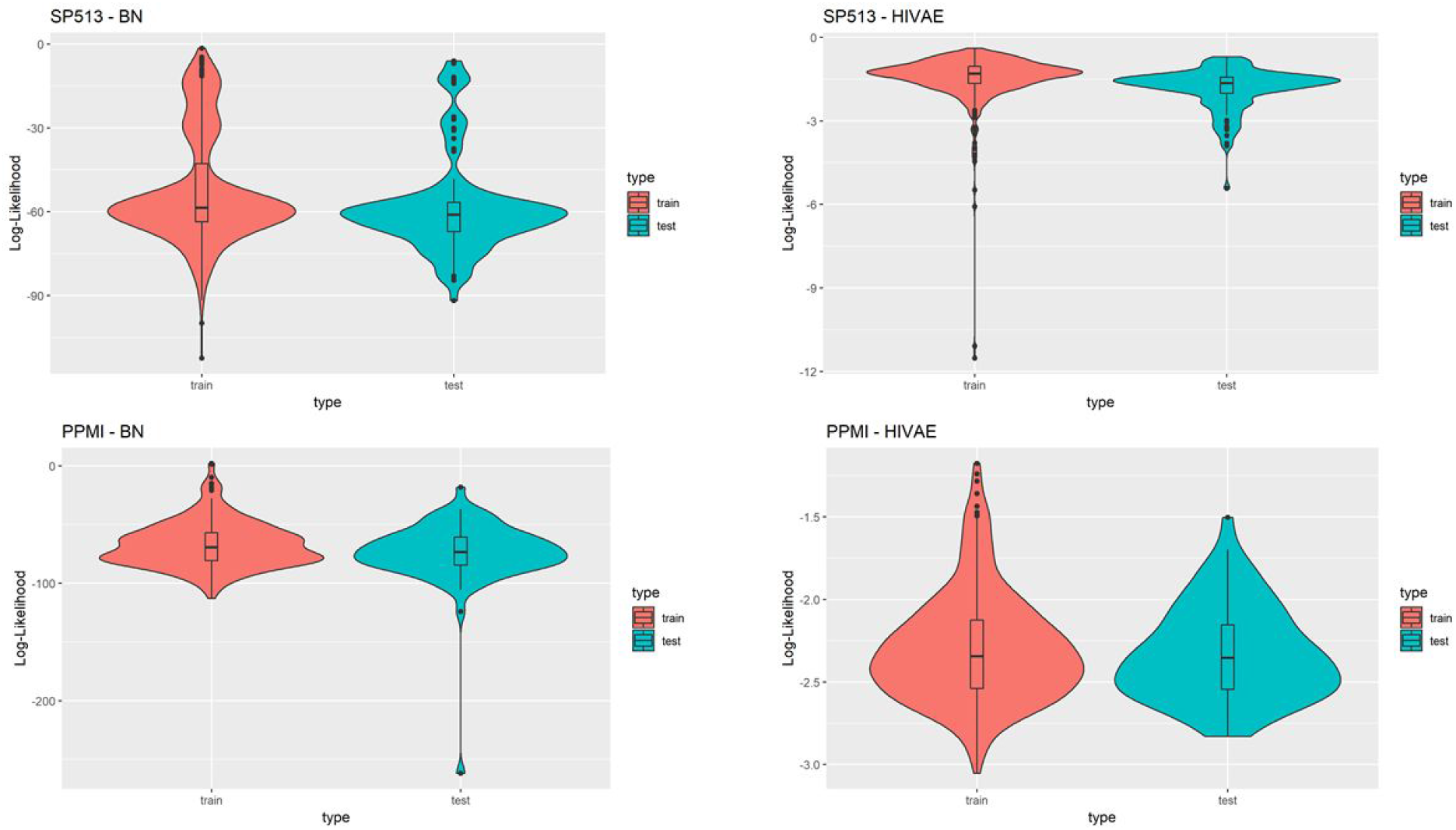
This figure compares the log-likelihoods of real patients in a training set (red) and a test set (blue) of the SP513 (top row) and PPMI datasets (bottom row) for the MBN and the HI-VAE models. The HI-VAE log-likelihoods are based on the participants included in the respective sets after averaging across all separate HI-VAE models (9 for SP513, 34 for PPMI).

### Simulation of Counterfactual Scenarios Match Expectations

Due to its nature as a hybrid of a BN and a generative neural network VAMBN allows for simulation of counterfactual scenarios via the “do” calculus, as explained in the Methods part. Figure 7a) demonstrates the effect of counterfactually altering UPDRS2 and UPDRS3 baseline scores of all patients in SP513 to the mean observed in PPMI, i.e. towards lower disease severity. As expected this resulted into a likewise shift of UPDRS3 scores (reflecting motor symptoms) at end of study.

**Figure 7.**
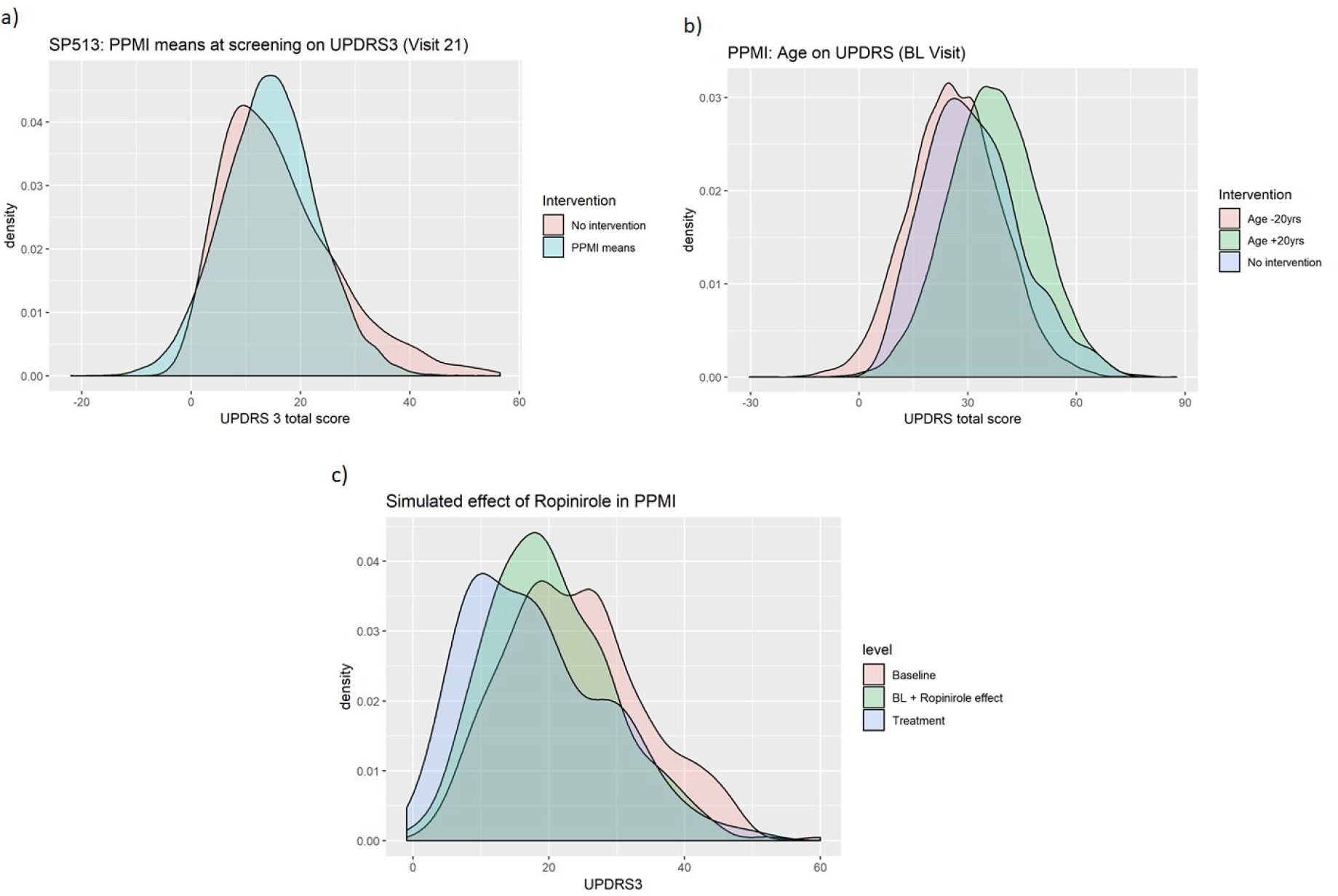
Counterfactual simulation of (a) lower disease severity in SP513 (shift of UPDRS3 scores at baseline to mean observed in PPMI at baseline); (b) shift of age 20 years up and down; (c) treatment effect of ropinirole in PPMI.

In PPMI making all patients 20 years younger shifts the distribution of UPDRS3 scores to the left (fewer motor symptoms), whereas making them 20 years older has the opposite effect (Figure 7b). Again, this effect matches expectations.

As a final example we demonstrate the possibility to counterfactually add a feature to PPMI via the approach described in the Methods part: We used the VAMBN model for SP513 to simulate the shift of the UPDRS3 scores at visit 15 under ropinirole treatment conditional on age, gender, height, weight as well as UPDRS2 and UPDRS3 baseline scores of patients observed in PPMI. That means there was only a simulated intervention into these features. By subtracting the simulated shift from the observed UPDRS3 off scores in PPMI we obtained a counterfactual treatment simulation with ropinirole. Figure 7c compares the observed UPDRS3 off and on scores (under L-DOPA treatment) to those simulated by VAMBN for ropinirole treatment. Further plots showing the effect at different PPMI visits are shown in section D of the Supplementary Materials. As expected, UPDRS3 scores simulated for ropinirole treatment are significantly shifted compared to observed UPDRS3 off scores, but are slightly higher than UPDRS3 on scores under L-DOPA. Indeed it has been suggested that efficacy of ropinirole is slightly lower than that of L-DOPA (Zhuo et al., 2017).

Overall these counterfactual simulations exemplify the possibilities of VAMBN and at the same time reconfirm that the model has learned the expected variable dependencies from data, because the simulation effects match expectations.

### Differential Privacy Respecting Modeling Training

As a last point we investigated differential privacy respecting model training of VAMBN. As indicated in the Methods part this can be realized by defining a certain privacy loss via constants (ε, δ) for each HI-VAE model trained within VAMBN. Smaller values for these constants generally impose stronger privacy guarantees, but make model training harder. To investigate this effect more quantitatively, Figures 7 shows the reconstruction errors of the HI-VAE models for the SP513 laboratory data at the first visit as a function of number of training epochs and in dependency on different values for ε, δ. For similar figures of the other visits, see section E of the Supplementary Materials. It can be observed that in dependency on these constants longer trainings and more data are required to achieve the same level of reconstruction error than for conventional model training without differential privacy.

**Figure 8.**
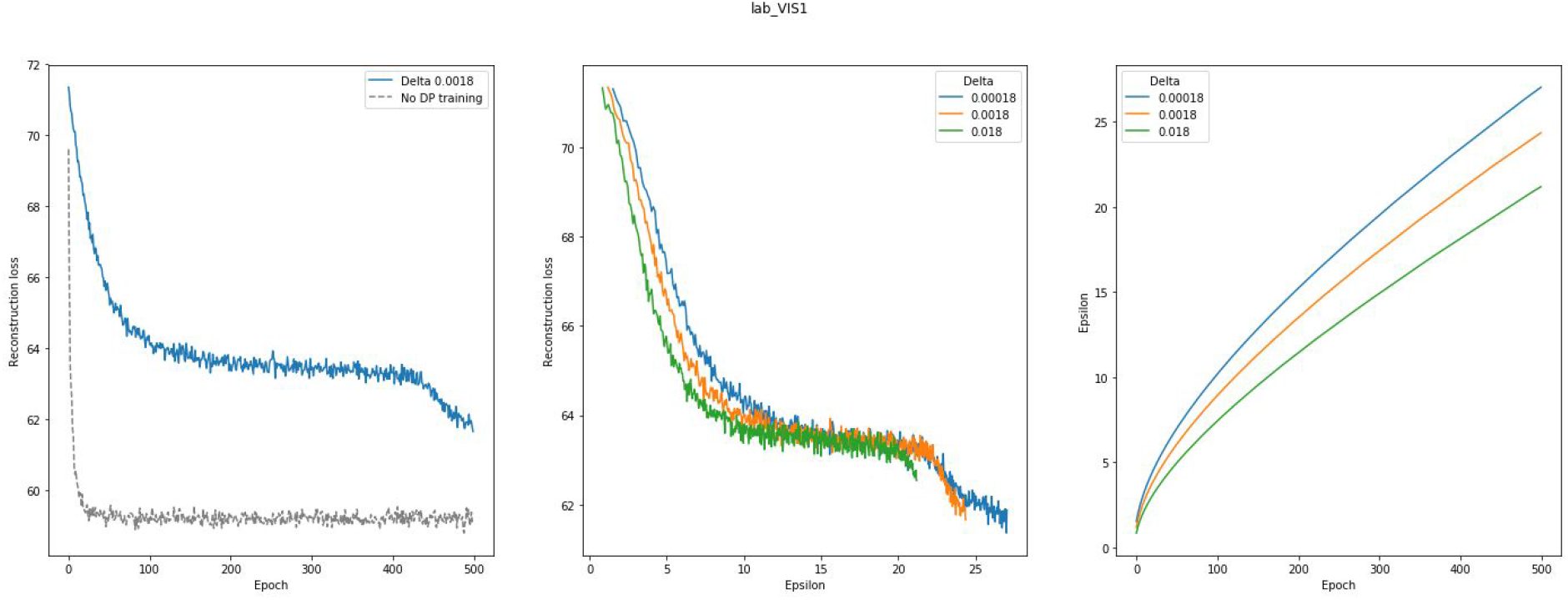
This figure shows the effects of differentially privacy respecting (DP) HI-VAE training on the HI-VAE step of the model. Left panel: Reconstruction loss change between DP and conventional model training for laboratory data at visit 1 for the SP513 study; middle panel: Epsilon plotted against reconstruction loss for different delta values; right panel: Epsilon over 500 epochs, given different deltas. A noise multiplier of 1.1, norm clipping at 1.6 and a learning rate of 0.01 were used. Further plots can be found in section E of the Supplementary Materials.

## Conclusion

Sensitive patient data require high standards for protection, as e.g. reinforced by the European Union through the General Data Protection Regulation (GDPR - https://eur-lex.europa.eu/eli/reg/2016/679/oj). However, at the same time these data are instrumental for biomedical research in the entire healthcare sector. Establishing a mechanism for sharing data across organizations without violating data privacy is therefore of utmost relevance for scientific progress. In this paper we build on the idea of developing generative models to simulate virtual patients based on data from clinical studies. A recent publication proposed to train Generative Adversarial Networks (GANs) based on few variables recorded from more than 6000 patients in the Systolic Blood Pressure Trial (Beaulieu-Jones et al., 2018). In contrast, our work focuses on the realistic situation regarding much smaller sample size coupled with significantly higher number of variables, which is common in many other medical fields, such as neurology. Further distinction points of our VAMBN method include the explicit modeling of time dependencies, as well as missing and heterogenous data. Moreover, VAMBN models can be interpreted via the MBN structure. As demonstrated in this work Bayesian Networks also open the door to simulating counterfactual scenarios within a well-established theoretical framework, which could e.g. help in the design of clinical trials.

Our results demonstrate that VAMBN models generally do not overfit and allow for a sufficiently realistic simulation of virtual patients. Hence, we see virtual patients as valuable for data mining purposes and for counterfactually merging different datasets. In addition, we demonstrated that data privacy respecting model training is in principle possible with VAMBN.

Of course, our work is not without limitations: Building VAMBN models requires (in contrast to GANs) a relatively detailed understanding of data and careful handling of missing values in particular. Moreover, VAMBN implies to train multiple neural networks, which usually requires a modern parallel computing architecture. It thus remains a subject of future research to investigate how VAMBN models could be made better accessible to practitioners in order to facilitate their use in a widespread manner. In the meantime we have made our python and R code available as part of the Supplementary material.

Overall we see our work as a useful complement to federated machine learning techniques, which together with virtual patient simulation tools could help to break data silos and thus enhance progress in biomedical research.

## Acknowledgements

Data used in the preparation of this article were obtained from the Parkinson’s Progression Markers Initiative (PPMI) database (www.ppmi-info.org/data). For up-to-date information on the study, visit www.ppmi-info.org. PPMI – a public-private partnership – is funded by the Michael J. Fox Foundation for Parkinson’s Research and funding partners. A list of names of all of the PPMI funding partners can be found at www.ppmi-info.org/about-ppmi/who-we-are/study-sponsors/.

## Abbreviations

VAMBN: Variational Autoencoder Modular Bayesian Network
BN: Bayesian Network
MBN: Modular Bayesian Network
VAE: Variational Autoencoder
HI-VAE: Heterogenous and Incomplete Data Variational Autoencoder
DAG: directed acyclic graph
MCAR: missing completely at random
MAR: missing at random
MNAR: missing not at random
BIC: Bayesian Information Score
PPMI: Parkinson’s Progression Marker Initiative
PD: Parkinson’s Disease
UPDRS: Unified Parkinson’s Disease Rating Scales
ESS: Epworth Sleepiness Scale
RBD: REM sleep behavior disorder
CSF: cerebrospinal fluid

