## Supplementary Materials for "Variational Autoencoder Modular Bayesian Networks (VAMBN) for Simulation of Heterogeneous Clinical Study Data"

The R and python code of the model can be found under https://github.com/elg34/VAMBN.

#### A. Iterative Training

The following plot contrast the log-likelihoods of real patients after the initial / base training of VAMBN (red) and one further iteration of the entire VAMBN training (consisting of a continued training of all HI-VAE models with a modified prior and re-estimation of the MBN, blue). Log-likelihoods shown for HI-VAE models are averaged over all modules (9 for SP513, 34 for PPMI).

### SP513

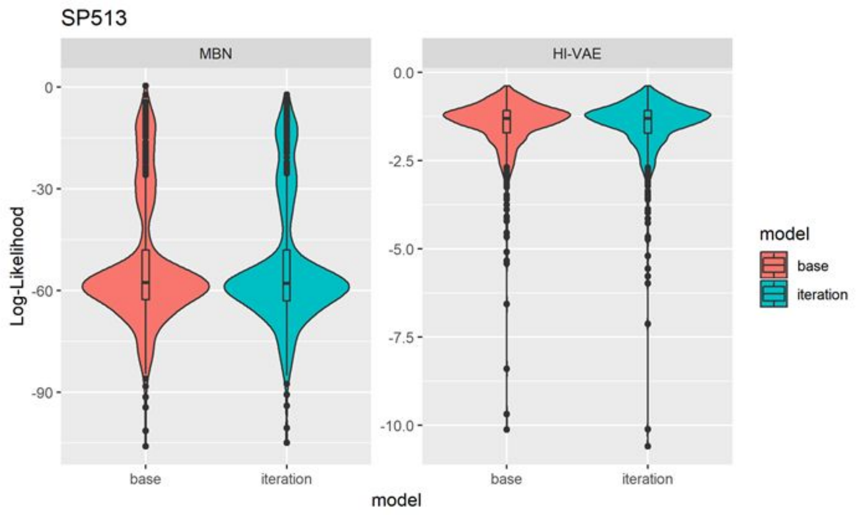

### PPMI

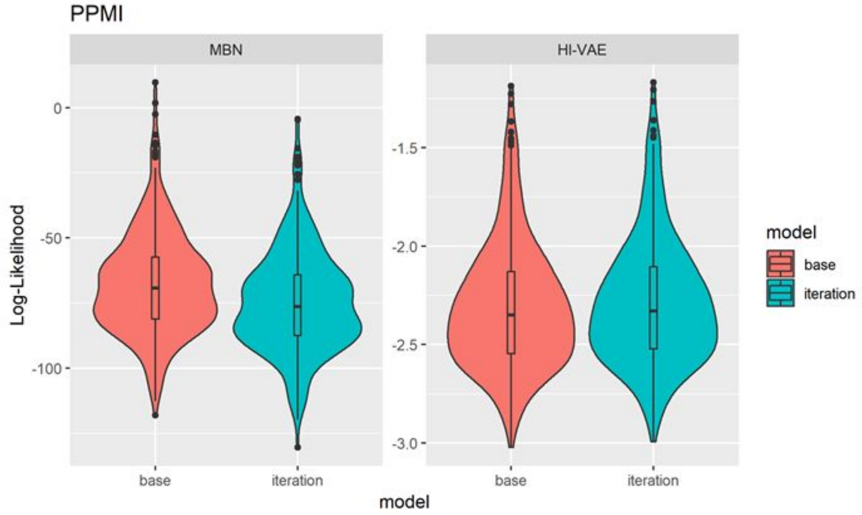


### B. Marginal Distributions: Simulated vs Real

This section shows additional marginal plots to the ones included in the main text. For the full set of marginal distributions, see [https://osf.io/qnemd/](https://osf.io/qnemd/).

## SP513

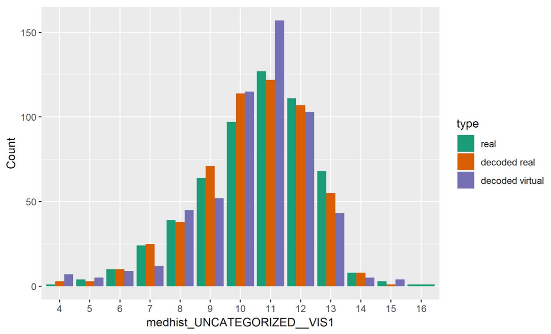


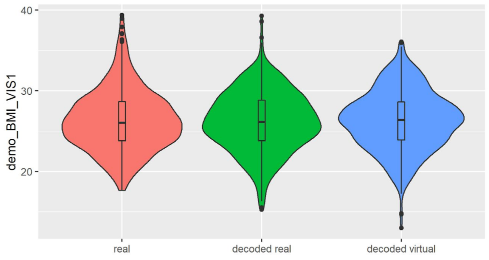


| Type | Mean | SD | 25% | Median | 75% |
|---|---|---|---|---|---|
| *real* | *26.34* | *3.81* | *23.81* | *26.03* | *28.67* |
| *decoded real* | *26.21* | *3.72* | *23.82* | *26.14* | *28.84* |
| *decoded virtual* | *26.34* | *3.68* | *23.91* | *26.38* | *28.64* |

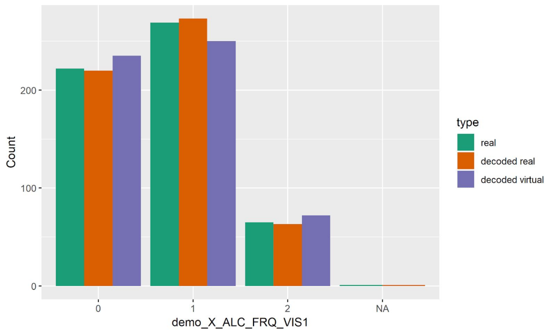


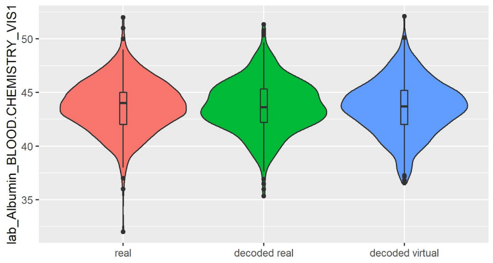


| Type | Mean | SD | 25% | Median | 75% |
|---|---|---|---|---|---|
| *real* | *43.77* | *2.38* | *42* | *44* | *45* |
| *decoded real* | *43.71* | *2.41* | *42.21* | *43.62* | *45.32* |
| *decoded virtual* | *43.61* | *2.51* | *42.02* | *43.7* | *45.19* |

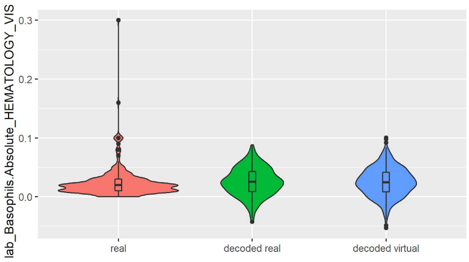


| Type | Mean | SD | 25% | Median | 75% |
|---|---|---|---|---|---|
| *real* | *0.02* | *0.02* | *0.01* | *0.02* | *0.03* |
| *decoded real* | *0.03* | *0.02* | *0.01* | *0.03* | *0.04* |
| *decoded virtual* | *0.03* | *0.02* | *0.01* | *0.02* | *0.04* |

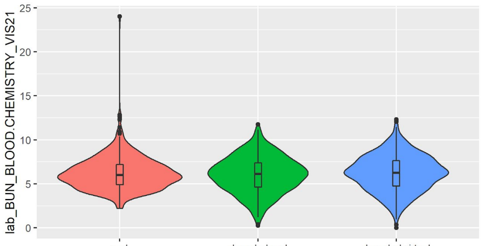


| Type | Mean | SD | 25% | Median | 75% |
|---|---|---|---|---|---|
| *real* | *6.24* | *1.95* | *4.9* | *6* | *7.2* |
| *decoded real* | *6.03* | *2.05* | *4.62* | *6.13* | *7.39* |
| *decoded virtual* | *6.22* | *2.04* | *4.76* | *6.26* | *7.63* |

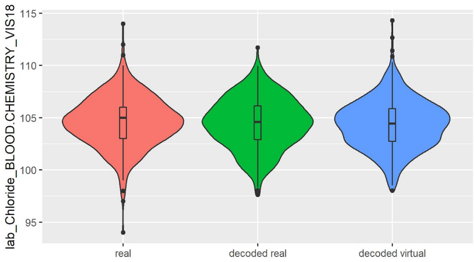


| Type | Mean | SD | 25% | Median | 75% |
|---|---|---|---|---|---|
| *real* | *104.6* | *2.42* | *103* | *105* | *106* |
| *decoded real* | *104.47* | *2.43* | *102.9* | *104.6* | *106.14* |
| *decoded virtual* | *104.38* | *2.35* | *102.74* | *104.43* | *105.87* |

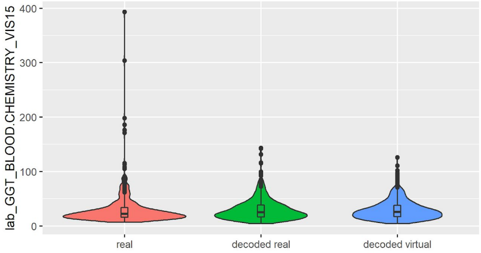


| Type | Mean | SD | 25% | Median | 75% |
|---|---|---|---|---|---|
| *real* | *30.75* | *31.46* | *16* | *22* | *34* |
| *decoded real* | *30.53* | *20.32* | *16.49* | *25.09* | *38.56* |
| *decoded virtual* | *29.73* | *18.32* | *16.86* | *25.7* | *37.9* |

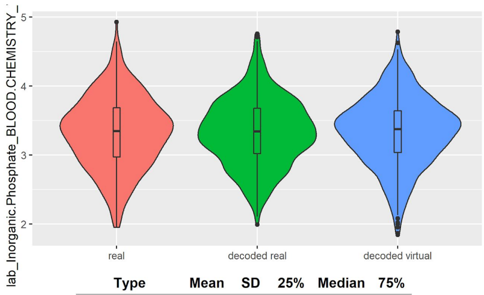


| Type | Mean | SD | 25% | Median | 75% |
|---|---|---|---|---|---|
| *real* | *3.33* | *0.52* | *2.97* | *3.35* | *3.68* |
| *decoded real* | *3.33* | *0.48* | *3.02* | *3.34* | *3.68* |
| *decoded virtual* | *3.34* | *0.5* | *3.03* | *3.37* | *3.64* |

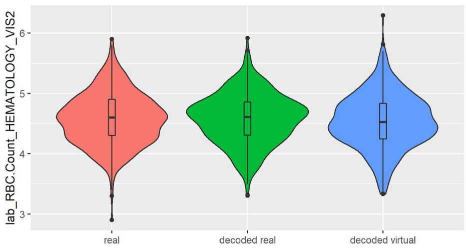


| Type | Mean | SD | 25% | Median | 75% |
|---|---|---|---|---|---|
| *real* | *4.57* | *0.43* | *4.3* | *4.6* | *4.9* |
| *decoded real* | *4.58* | *0.42* | *4.3* | *4.61* | *4.86* |
| *decoded virtual* | *4.54* | *0.45* | *4.25* | *4.52* | *4.83* |

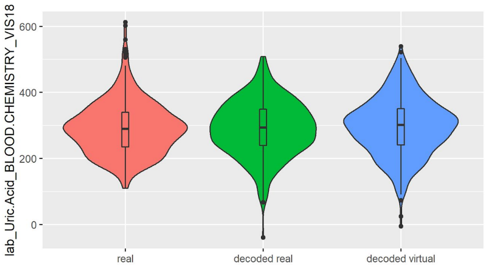


| Type | Mean | SD | 25% | Median | 75% |
|---|---|---|---|---|---|
| *real* | *292.41* | *79.71* | *235* | *290* | *340* |
| *decoded real* | *291.23* | *83.29* | *239.07* | *294.22* | *349.53* |
| *decoded virtual* | *297.36* | *86.55* | *241.25* | *301.37* | *351.04* |

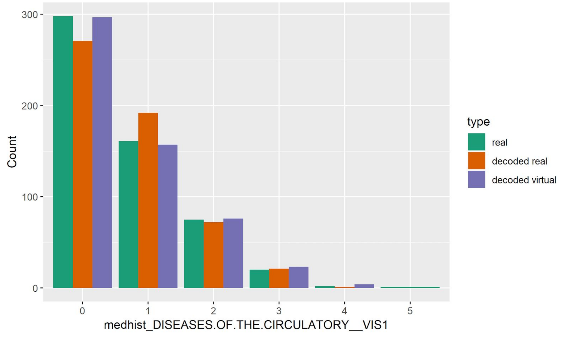

### PPMI

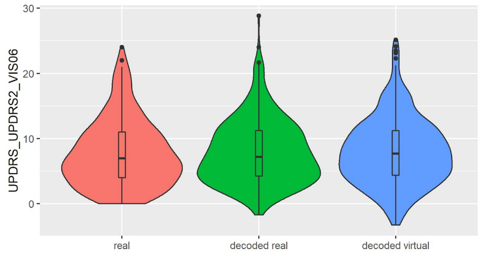


| Type | Mean | SD | 25% | Median | 75% |
|---|---|---|---|---|---|
| real | 7.73 | 5.12 | 4 | 7 | 11 |
| decoded real | 7.94 | 4.91 | 4.27 | 7.18 | 11.21 |
| decoded virtual | 8 | 5.14 | 4.39 | 7.68 | 11.21 |

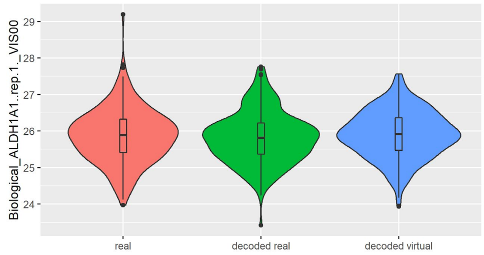


| Type | Mean | SD | 25% | Median | 75% |
|---|---|---|---|---|---|
| real | 25.87 | 0.74 | 25.42 | 25.89 | 26.33 |
| decoded real | 25.82 | 0.71 | 25.37 | 25.81 | 26.22 |
| decoded virtual | 25.91 | 0.65 | 25.47 | 25.92 | 26.37 |

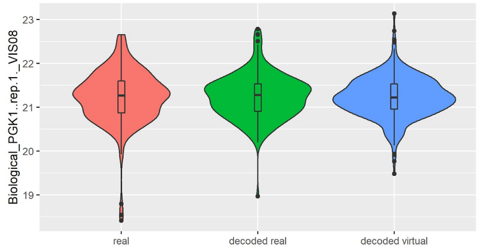


| Type | Mean | SD | 25% | Median | 75% |
|---|---|---|---|---|---|
| real | 21.23 | 0.61 | 20.87 | 21.27 | 21.6 |
| decoded real | 21.24 | 0.5 | 20.91 | 21.28 | 21.53 |
| decoded virtual | 21.23 | 0.45 | 20.96 | 21.22 | 21.53 |

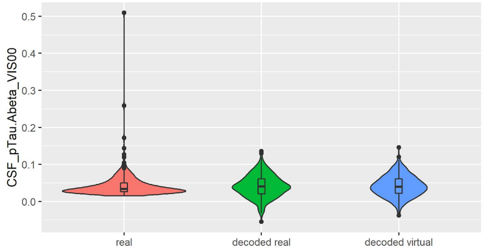


| Type | Mean | SD | 25% | Median | 75% |
|---|---|---|---|---|---|
| real | 0.04 | 0.03 | 0.03 | 0.03 | 0.05 |
| decoded real | 0.04 | 0.03 | 0.02 | 0.04 | 0.06 |
| decoded virtual | 0.04 | 0.03 | 0.02 | 0.04 | 0.06 |

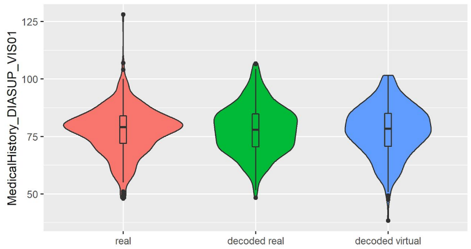


| Type | Mean | SD | 25% | Median | 75% |
|---|---|---|---|---|---|
| real | 77.56 | 10.25 | 72 | 79 | 84 |
| decoded real | 78.06 | 10.51 | 70.49 | 77.83 | 84.75 |
| decoded virtual | 77.73 | 10.92 | 70.78 | 78.3 | 85.02 |

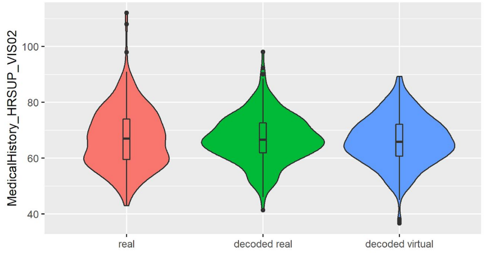


| Type | Mean | SD | 25% | Median | 75% |
|---|---|---|---|---|---|
| real | 67.1 | 10.78 | 59.5 | 67 | 74 |
| decoded real | 67 | 8.61 | 61.88 | 66.61 | 72.66 |
| decoded virtual | 66.17 | 8.79 | 60.67 | 65.82 | 72.17 |

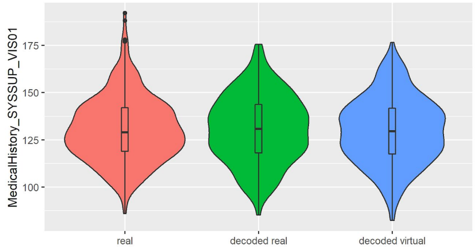


| Type | Mean | SD | 25% | Median | 75% |
|---|---|---|---|---|---|
| real | 130.43 | 17.04 | 119 | 129 | 142 |
| decoded real | 130.87 | 17.9 | 118.11 | 130.87 | 143.81 |
| decoded virtual | 130.01 | 17.57 | 117.46 | 129.53 | 141.76 |

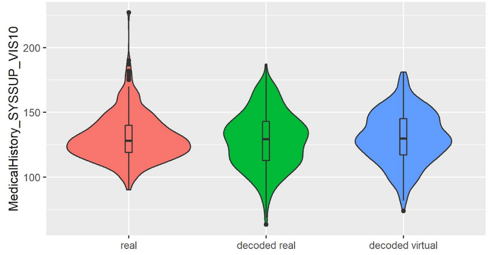


| Type | Mean | SD | 25% | Median | 75% |
|---|---|---|---|---|---|
| real | 130.58 | 18 | 119 | 128 | 140 |
| decoded real | 128.53 | 21.41 | 112.77 | 129.19 | 143.03 |
| decoded virtual | 130.39 | 19.78 | 117.05 | 129.8 | 145.04 |

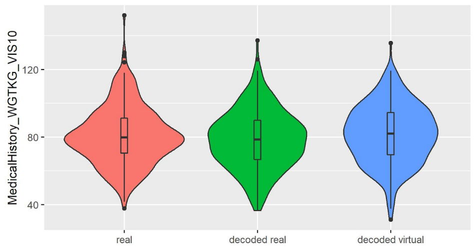


| Type | Mean | SD | 25% | Median | 75% |
|---|---|---|---|---|---|
| *real* | *80.75* | *17.07* | *70.5* | *79.8* | *91.2* |
| *decoded real* | *78.44* | *17.25* | *66.75* | *78.53* | *89.91* |
| *decoded virtual* | *81.77* | *17.31* | *69.56* | *82.06* | *94.52* |

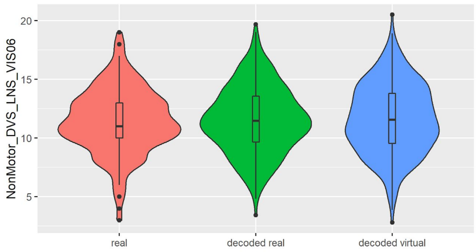


| Type | Mean | SD | 25% | Median | 75% |
|---|---|---|---|---|---|
| *real* | *11.45* | *2.78* | *10* | *11* | *13* |
| *decoded real* | *11.55* | *2.82* | *9.67* | *11.46* | *13.57* |
| *decoded virtual* | *11.63* | *2.98* | *9.54* | *11.57* | *13.8* |

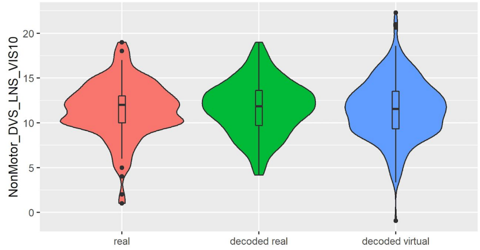


| Type | Mean | SD | 25% | Median | 75% |
|---|---|---|---|---|---|
| *real* | *11.5* | *3.1* | *10* | *12* | *13* |
| *decoded real* | *11.71* | *2.97* | *9.69* | *11.84* | *13.61* |
| *decoded virtual* | *11.52* | *3.25* | *9.33* | *11.55* | *13.51* |

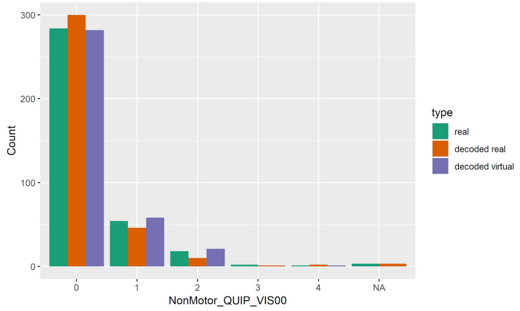


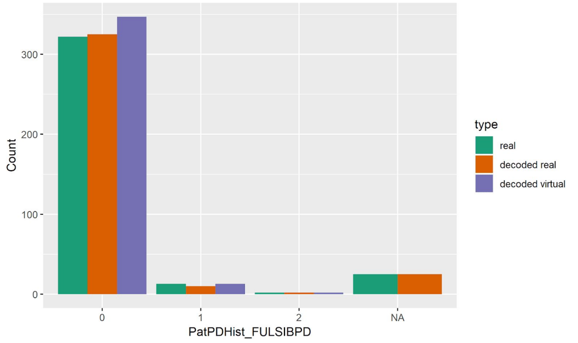


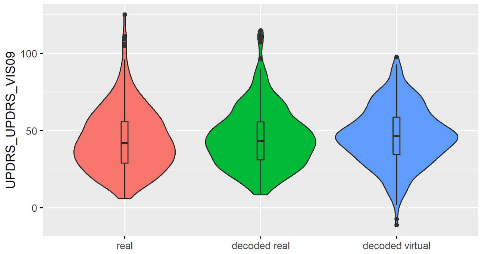


| Type | Mean | SD | 25% | Median | 75% |
|---|---|---|---|---|---|
| *real* | *43.98* | *20.08* | *29* | *42* | *56* |
| *decoded real* | *45.22* | *19.28* | *31.15* | *43.13* | *55.65* |
| *decoded virtual* | *46.43* | *18.48* | *34.66* | *46.3* | *58.59* |

### C. Simulated Patients: Group Comparisons at Further Visits

## SP513

Distribution of UPDRS3 scores in SP513. The plot depicts in red UPDRS3 scores of real SP513 patients under placebo and ropinirole at visits 18 and 21. In blue the distribution of the UPDRS3 score in the same number of virtual patients is shown. Effect sizes and corresponding p-values obtained from two separate one-way ANOVAs comparing placebo and drug treatment for real and virtual patients are shown in the table at the bottom.

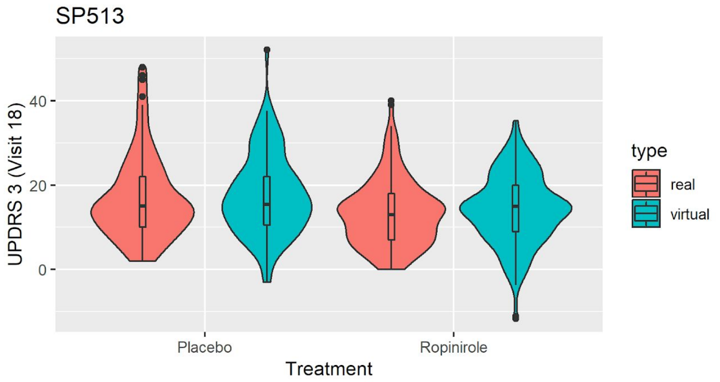


| Type | p value | ges |
|---|---|---|
| *real* | *0.001* | *0.04* |
| *virtual* | *0.002* | *0.03* |

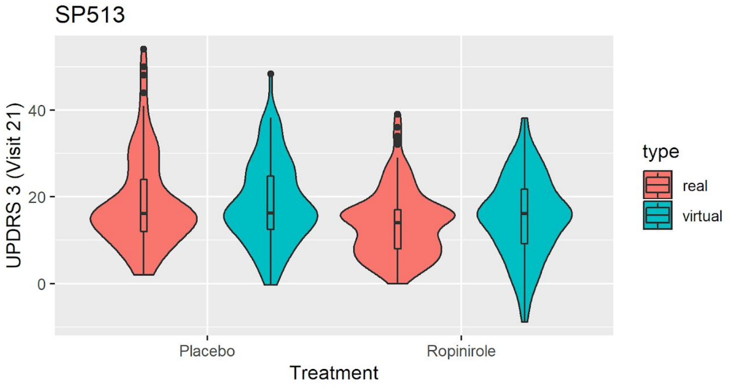


| Type | p value | ges |
|---|---|---|
| *real* | *<.001* | *0.06* |
| *virtual* | *0.005* | *0.03* |

### PPMI

Distribution of original (purple) and decoded (red) UPDRS3 scores of real PPMI de novo PD patients at visit 6 and 8 in comparison to PPMI healthy controls (blue). UPDRS3 scores of virtual PD patients are shown in green. The table at the bottom shows differences in UPDRS3 scores between original PD, decoded real PD and virtual PD patients compared to PPMI healthy controls, showing p-value and effect size from Mann-Whitney U tests.

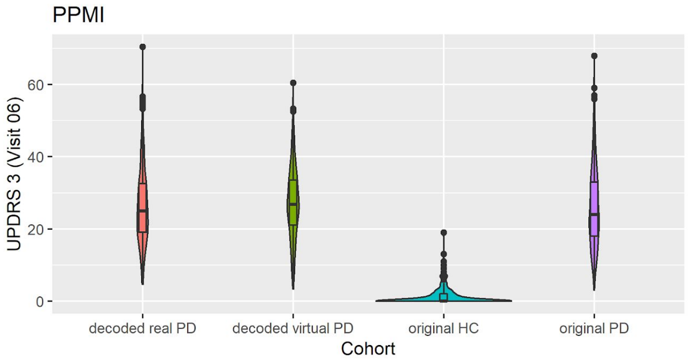


| Type | p value | r |
|---|---|---|
| *real PD- HC* | *<0.001* | *0.772* |
| *decoded real PD - HC* | *<0.001* | *0.794* |
| *decoded virtual PD - HC* | *<0.001* | *0.793* |

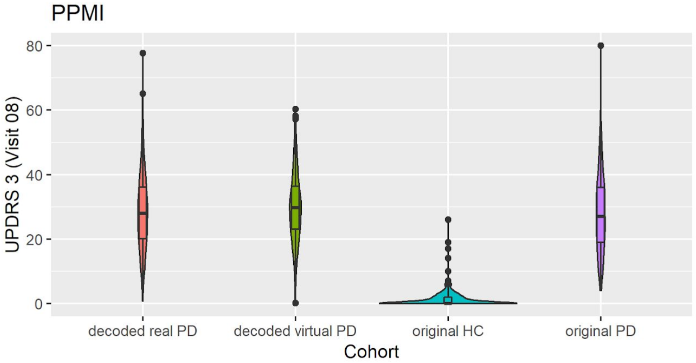


| Type | p value | r |
|---|---|---|
| *real PD- HC* | *<0.001* | *0.762* |
| *decoded real PD - HC* | *<0.001* | *0.779* |
| *decoded virtual PD - HC* | *<0.001* | *0.783* |

### D. Counterfactual Simulations: Further Visits

In addition to the results in Figure 7 (main text), the following figures show the same results for different visits (06,08,10) in the PPMI study.

Simulated effect of Ropinirole in PPMI (V06)

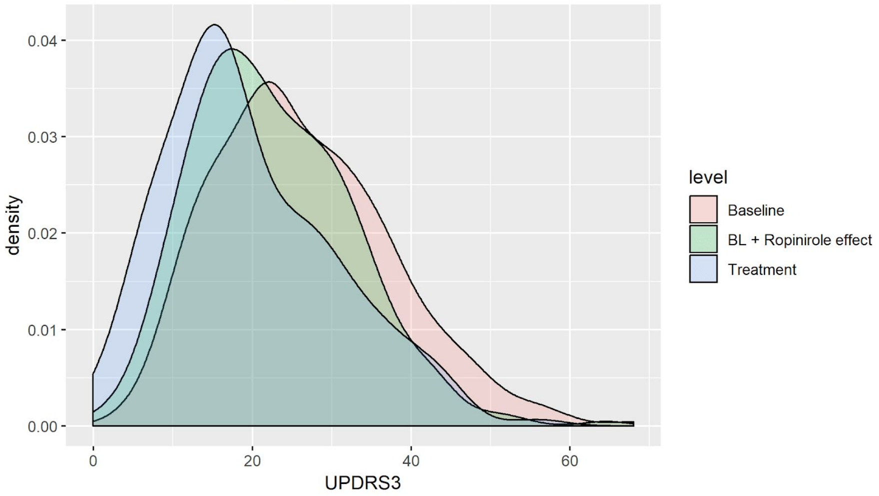


Simulated effect of Ropinirole in PPMI (V08)

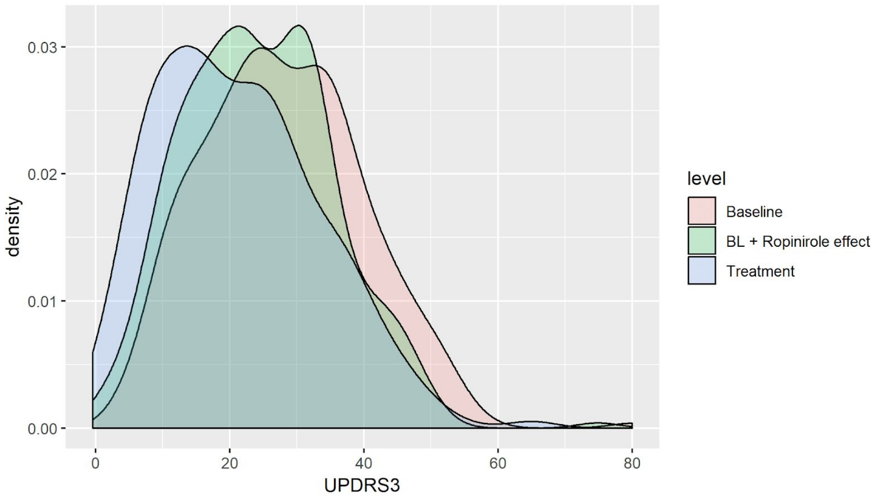


Simulated effect of Ropinirole in PPMI (V10)

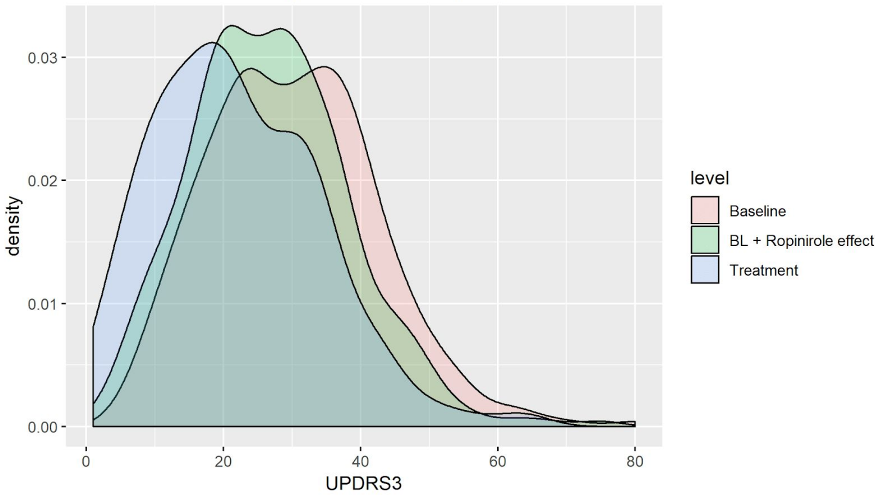

### E. Differential Privacy Respecting Modeling Training

Below are the results for the differentially private training of all SP513 laboratory data (5 visits). The leftmost panel shows the change in reconstruction loss, the middle panel shows the reconstruction loss across different values of epsilon for different deltas and the rightmost panel shows the relationship between the amount of training received and epsilon/delta values. For all visits, we used a noise multiplier of 1.1, norm clipping at 1.6 and a learning rate of 0.01.

lab_VIS1

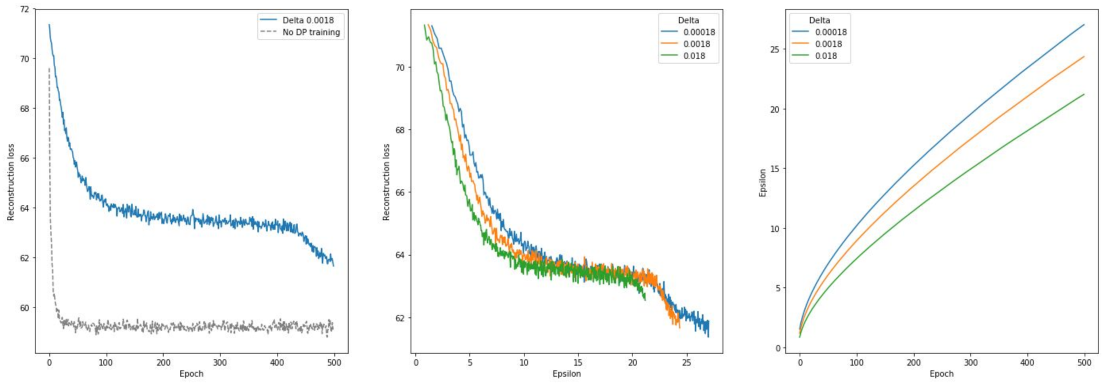


lab_VIS2

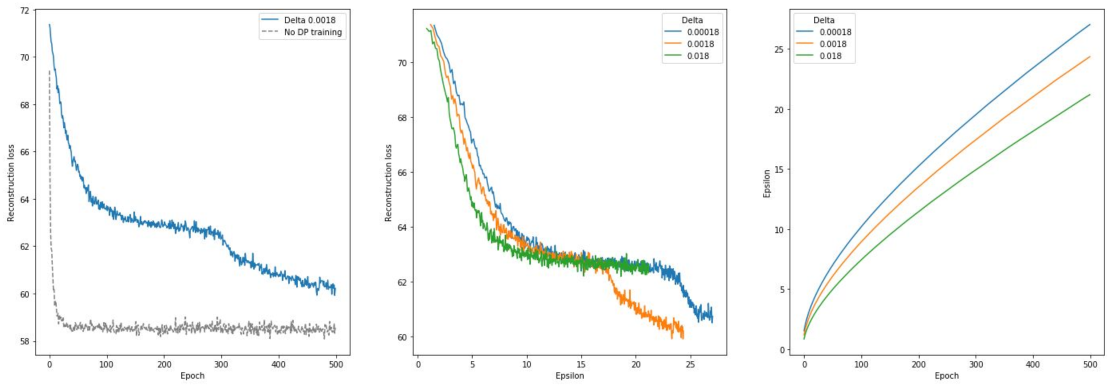

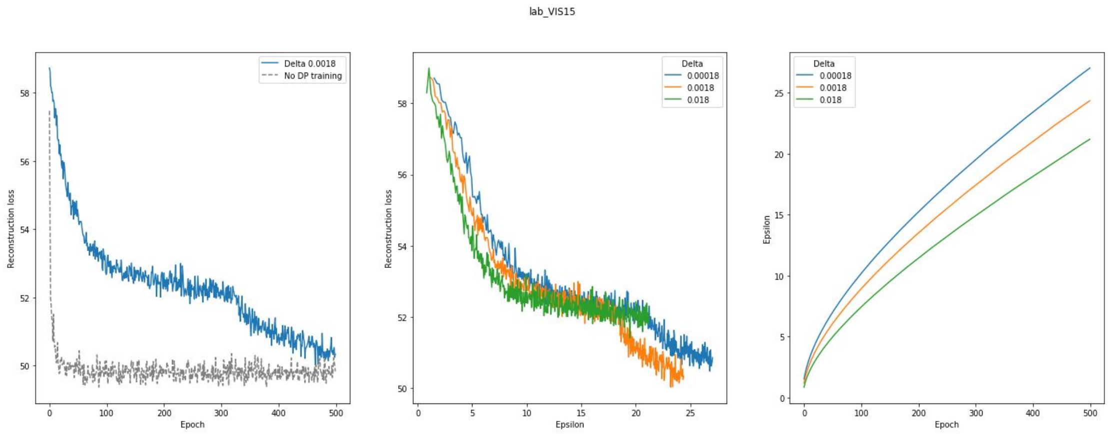
lab_VIS15
Delta 0.0018
No DP training
Reconstruction loss
Epoch
Delta
0.00018
0.0018
0.018
Reconstruction loss
Epsilon
Delta
0.00018
0.0018
0.018
Epsilon
Epoch

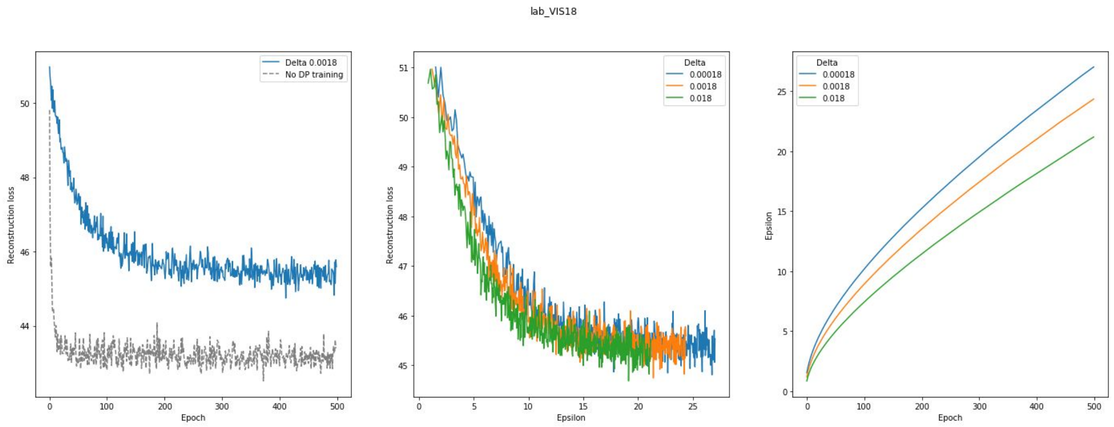
lab_VIS18
Delta 0.0018
No DP training
Reconstruction loss
Epoch
Delta
0.00018
0.0018
0.018
Reconstruction loss
Epsilon
Delta
0.00018
0.0018
0.018
Epsilon
Epoch

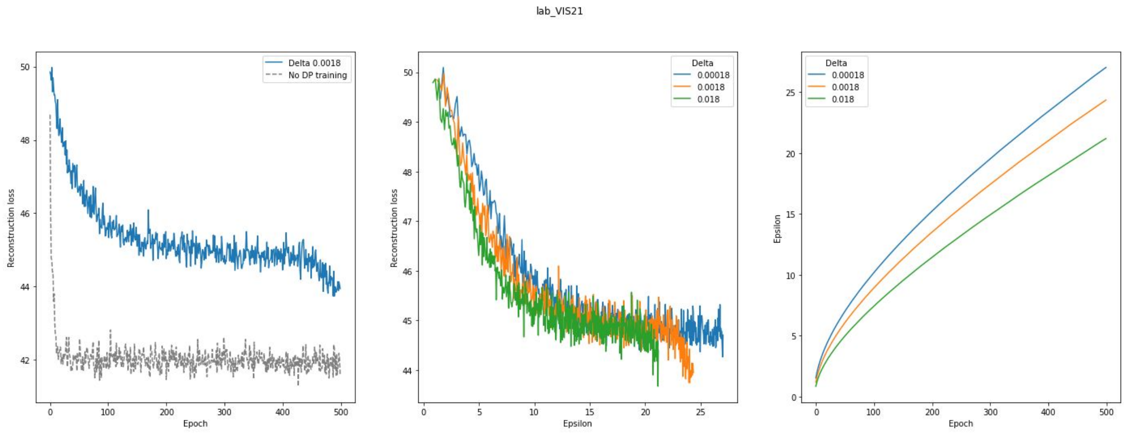
lab_VIS21
Delta 0.0018
No DP training
Reconstruction loss
Epoch
Delta
0.00018
0.0018
0.018
Reconstruction loss
Epsilon
Delta
0.00018
0.0018
0.018
Epsilon
Epoch
